# Trait-dependent biogeography offers insights on the dispersal of *Meiogyne* (Annonaceae) across the Australasia-Pacific region

**DOI:** 10.1101/2024.09.19.614018

**Authors:** Ming-Fai Liu, Jérôme Munzinger, Piya Chalermglin, Junhao Chen, Bine Xue, Richard M. K. Saunders

## Abstract

*Meiogyne* is a genus of trees and treelets occurring in Indomalaya and Australasia-Pacific, an unusually wide distribution across Australasia and Western Pacific compared to the rest of the family Annonaceae. Previous chloroplast phylogenies of the genus offered poor resolution and support for many internal nodes. Here, a molecular phylogeny was reconstructed based on seven chloroplast and 11 nuclear markers of 33 *Meiogyne* taxa (*ca.* 70% sampling). The combined dataset generated a well resolved and supported phylogeny. Estimation of divergence time was calibrated with two fossils using uncorrelated lognormal relaxed clock model. Trait-dependent and trait-independent biogeographical models in BioGeoBEARS were compared using AICc weight and likelihood ratio test. The results suggest that narrow monocarp width and fruit colour associated with bird dispersal are correlated with increased macroevolutionary dispersal. Under the best-fitting monocarp width-dependent DEC model, a single colonisation event from Sunda to Sahul during the middle to late Miocene and two dispersal events from New Guinea and Australia into the Pacific during the late Miocene to early Pliocene were detected. BayesTraits analysis strongly supports a correlation between narrow fruits and avian fruit colours. This study reveals that *Meiogyne* lineages with narrow fruitlets and fruit colour associated to bird dispersal (black, red & orange) are associated with increased macroevolutionary dispersal. Bird dispersal and the associated traits may be important drivers for macroevolutionary dispersal for plants with fleshy fruits in Australasia-Pacific.

## 1 INTRODUCTION

The functional traits of diaspores can have a major influence on the dispersal range and fitness of a species, with diaspore traits often associated with the type of dispersal agents (Fischer and Chapman, 1993; Voigt et al., 2004; Flörchinger et al., 2010; Donatti et al., 2011; Schaefer and Ruxton, 2011; Valido et al., 2011; Hodgkison, 2013), which in turn affect the dispersal distance and clustering of diaspores (Wenny and Levey, 1998; Seidler and Plotkin, 2006; Jordano et al., 2007; Nathan et al., 2012). It is perhaps not surprising that diaspore traits have been hypothesised to affect long-distance dispersal (LDD) or macroevolutionary dispersal (Darwin, 1859; Nathan, 2006; Losos and Ricklefs, 2009). Due to the rarity and stochastic nature of LDD, direct data collection and analysis of LDD are usually difficult (Cain et al., 2000; Nathan et al., 2003; Nathan, 2006) and statistical biogeographical models that retrospectively assess LDD often lack the capacity to take biological factors into account (Matzke, 2013; Sukumaran and Knowles, 2018). Recent advances in trait-incorporated biogeographical models, however, have enabled the testing of trait-related biogeographical hypotheses (MatosDMaraví et al., 2018; Blom et al., 2019; Lu et al., 2019; Klaus and Matzke, 2020).

Many tropical rainforest tree species rely on endozoochory for diaspore dispersal (Howe and Smallwood, 1982), in which frugivores are rewarded with nutrients in exchange for displacing diaspores from the parent plants. Diaspore traits are known to influence frugivore assemblages as a result of diffuse co-evolution (Gautier-Hion et al., 1985; Voigt et al., 2004; Shanahan et al., 2001; Lomáscolo et al., 2008). Because of their fast and directed movement, frugivorous migratory birds can disperse seeds further from the parent plants compared to non-migratory animals, and this is likely to promote LDD across the evolutionary timescale (Nathan, 2006; Nathan et al., 2008). Due to limited dexterity and gape size, birds are largely restricted to the consumption of fruits narrower than 2.2 cm, although hornbills and some fruit pigeons can eat larger fruits (Leighton and Leighton, 1983; Kitamura, 2011). The diets of frugivorous birds consist of fruits with a broad colour range, including black, red, orange, yellow, purple, white, blue and pink, but rarely those that are dull coloured (Janson, 1983; Knight and Siegfried, 1983; Willson et al., 1990; Voigt et al., 2004). Fruit colour and size have been implicated in the macroevolutionary dispersal of plants within the framework of molecular phylogenies (Lu et al., 2019; Onstein et al., 2019; Klaus and Matzke, 2020). In this study, we test the effect of two fruit traits related to bird dispersal—fruit size and colour—on macroevolutionary dispersal of a tropical woody lineage using trait-dependent biogeographic models.

The pantropical family Annonaceae (∼2,500 spp.) is restricted to mostly tropical rainforests and relies almost exclusively on fruits with fleshy pulp for frugivorous dispersal (Van Setten and Koek-Noorman, 1992; Onstein et al., 2019). With few exceptions, Annonaceae fruits are generally apocarpous, comprising multiple fruitlets (‘monocarps’, which are borne on the receptacle), with each monocarp derived from a separate, unfused carpel within the flower. The diversity of fruit traits and associated animal dispersal syndromes render this family ideal for testing trait-dependent macroevolutionary dispersal hypotheses. In a previous family-wide biogeographical study (Onstein et al., 2019), multiple frugivore-related traits were shown to drive LDD. However, the effects of these traits appeared to be highly context-specific based on the type of disjunctions, suggesting bird and mammal dispersal agents might have influenced macroevolutionary dispersal of the plants differently in different palaeogeographical settings. The current study focuses on the genus *Meiogyne* (Annonaceae subfamily Malmeoideae tribe Miliuseae; Chatrou et al., 2012; Guo et al., 2017), which exhibits an unusually wide distribution and species richness across Australasia and Western Pacific compared to the rest of the family (Turner and Utteridge, 2017). Fruit morphology furthermore varies greatly among species and might have been the result of frugivore adaptation (Van Heusden, 1994; Thomas et al., 2012). Molecular phylogenetic reconstructions of *Meiogyne* based on chloroplast DNA (cpDNA) have retrieved a basal Indomalayan grade and an Australasia-Pacific Clade (APC), with most of the LDD events suggested to have occurred in the latter (Thomas et al., 2012; Xue et al., 2014). Nonetheless, the recalcitrant nature of the cpDNA phylogeny, especially in the nodes where disjunctions have occurred, prevented meaningful interpretation of historical biogeographical events.

In the present study, we aim to reconstruct a well resolved phylogeny, which will then serve as a basis to test the hypothesis that dispersal-related traits—fruit colour and monocarp width—affect macroevolutionary dispersal in *Meiogyne*. The trait-dependent model in BioGeoBEARS (Klaus and Matzke, 2020) allows estimation of binary character state transition rates and the relative influence of the binary character states on anagenetic dispersal, as well as assessment of migration routes used in the colonisation of Australasia-Pacific region through ancestral range reconstruction.

## 2 MATERIALS AND METHODS

### 2.1 Taxon and DNA sampling

For molecular phylogenetic reconstruction, 60 accessions were sampled. Thirty-four ingroup accessions were included, representing 27 of the 39 described species of *Meiogyne* (*ca.* 70% taxon sampling), of which two were newly sampled: *M. papuana* I.M.Turner & Utteridge and *M. punctulata* (Baill.) I.M.Turner & Utteridge (two accessions from New Caledonia, one representing the synonym *M. tiebaghiensis* (Däniker) Heusden). In addition, one undescribed species from New Guinea, one accession of *M. cylindrocarpa* (Burck) Heusden from Vanuatu (representing the synonym *Oncodostigma wilsonii* Guillaumin), one accession of *M. virgata* (Blume) Miq. from Thailand, and one accession of *M. virgata* from Vietnam (representing the synonym *M. monogyna* (Merr.) Bân) were also included. The sister relationship between *Meiogyne* and Sapranthinae have been reported in some studies (Chaowasku et al., 2014; Ortiz-Rodriguez et al., 2016; Xue et al., 2018), although not always retrieved (Mols et al., 2004; Couvreur et al., 2019); a sister relationship between *Meiogyne* and the Sapranthinae-*Wuodendron* clade was later retrieved (Xue et al, 2020). A total of 26 outgroup taxa from tribes Miliuseae and Maasieae were therefore included. For molecular divergence time estimation, outgroup taxa were further supplemented with 74 species across the magnoliids, bringing the total number of accessions to 134.

Seven cpDNA regions (*matK*, *ndhF*, *ndhF*-*rpl32*, *rbcL*, *rpl32*-*trnL*, *trnL*-*F* and *ycf1*) and eleven newly sequenced low-copy nuclear gene intron regions (*ATPQ*, *COX6A*, *CHER1*, *DDB2*, *DJC65*, *EIF3K*, *MNJ7.16*, *NIA*, *PDF5*, *STG1* and *MHF15.12*) were used. The nDNA intron regions were amplified using available Annonaceae nuclear marker primers or primers newly designed by Primer3 (Geneious) at the flanking exon regions of the target low-copy nuclear genes present in *M. hainanensis* (Merr.) Bân leaf transcriptome (unpublished). Low-copy nuclear gene regions were randomly selected from the *M. hainanensis* leaf transcriptome if they met the following criteria: (1) no more than three copies exist in the genome of *Arabidopsis*, *Populis*, *Vitis* or *Oryza* (Duerte et al., 2010); (2) paralogs absent in the leaf transcriptome; and (3) paralogs absent in Polymerase Chain Reaction (PCR) products. Exon-intron boundaries were determined by aligning the transcripts of *M. hainanensis* with homologous transcripts and gene sequences in *Arabidopsis thaliana* (L.) Heynh. The list of primers used is summarised in Appendix S1. Due to cost constraints, nuclear markers were only amplified for the subfamily Malmeoideae.

### 2.2 DNA extraction, amplification and sequencing

DNeasy Plant Mini Kit (Qiagen, Hilden, Germany) and a modified cetyl trimethyl ammonium bromide (CTAB) method (Doyle and Doyle, 1987; Erkens et al., 2008) were used to extract total genomic DNA from herbarium or silica-dried leaf materials, with the latter used wherever possible. PCR mixtures were prepared as recommended by the GoTaq Flexi DNA polymerase kit protocol (Promega, Madison, Wisconsin, USA). The following temperature programme was used: denaturation at 95 °C for 3 min; 35 cycles of denaturation at 95 °C for 30 s; primer annealing at 49–60 °C (0–5 °C below the T_m_) for 30 s; and primer extension at 72 °C for 30–75 s, with final extension at 72 °C for 7 min. Potential paralogy or non-specific amplification were assessed under gel electrophoresis. Amplicons were subsequently sent for commercial Sanger sequencing (BGI Group, Hong Kong).

### 2.3 Alignment and phylogenetic analysis

Nucleotide sequences were assembled and aligned using the MAFFT plugin with default settings (Katoh et al., 2002) in Geneious v.7.0.3 (https://www.geneious.com) before manual optimisation. A 6-bp inversion present in the *ndhF*–*rpl32* spacer was reverse-complemented, and polyA/T in *rpl32-trnL* and *ndhF-rpl32* spacers and the *trnL-F* region were excluded, as recommended by Thomas et al. (2012). Sequences absent were treated as missing data.

Maximum parsimony (MP) analyses were undertaken using PAUP v.4.0b10 (Swofford, 2002). All characters were treated as independent and unordered, and weighted equally. A heuristic search was performed with an unlimited number of trees retained during searches, 10,000 random-addition-sequence replicates, tree bisection-reconnection (TBR) branch swapping, and an upper limit of 100 trees saved per replicate. The best trees were summarised as a strict consensus tree. Non-parametric bootstrapping was performed with 1,000 replicates following the above specifications but with 100 random-addition-sequence replicates.

Maximum likelihood (ML) analyses were performed using in RAxML v.8.2.12 (Stamatakis, 2014) provided by the CIPRES Science Gateway (Miller et al., 2010). To deduce the best-fitting molecular evolution models and the optimal data partitioning schemes, PartitionFinder v.2 (Lanfear et al., 2017) was used, with 18 cpDNA and nDNA regions predefined according to their region identity. Since RAxML only allows a single model of rate heterogeneity in partitioned analyses, separate PartitionFinder analyses were undertaken on the partitioned dataset for each type of rate heterogeneity: the general time-reversible model without rate heterogeneity (GTR) and GTR model with gamma distributed rates across sites (GTR+Γ). The GTR+Γ+I model was not included since it is not recommended by the RAxML developers. Partitioned analyses were then performed with the best molecular evolution model to estimate the best-fitting partition scheme for each dataset, as shown in Appendix S2. A total of 1000 ML trees were inferred from distinct random-addition-sequence MP starting trees. Non-parametric bootstrapping was subsequently performed with 1,000 iterations.

Bayesian inference (BI) was performed using MrBayes v.3.2.6 (Ronquist et al., 2012) under the NSF Extreme Science & Engineering Discovery Environment (XSEDE) provided by the CIPRES Science Gateway. Optimal substitution models and partitioning schemes (Appendix S3) were determined by PartitionFinder v.2 (Lanfear et al., 2017). Four independent runs of Metropolis-coupled Markov chain Monte Carlo (MCMCMC) analyses were run. Each independent run consisted of one cold and three incrementally heated Markov chains with a temperature parameter of 0.08, and was run for 20 million generations and sampled every 1,000th generation. To reduce the probability of entrapment in local tree length optima, the mean branch length prior was reset from the default mean (0.1) to 0.01 (brlenspr=unconstrained:exponential(100.0)) (Brown et al., 2010; Marshall, 2010). The standard deviation of split frequencies was examined with values < 0.005 indicating good convergence. Tracer v.1.7.1 (Rambaut et al., 2018) was used to assess whether the MCMC parameters were drawn from stationary and unimodal distributions, and if sufficient effective sample size for each parameter was attained (ESS > 200). Convergence of posterior probabilities of splits between different runs, stationarity of posterior probabilities of splits within runs, tree topology autocorrelation, and approximate-ESS of tree topologies were assessed using RWTY (Warren et al., 2017). The first 25% samples of each MCMCMC run were discarded as burn-in. The remaining post-burn-in samples were then used to compute a 50% majority-rule consensus tree.

Support for internal nodes was assessed using bootstrap support (BS) in the MP and ML trees, and posterior probability (PP) in the Bayesian tree (interpreted as follows: 0–50% BS = not supported; 50–74% BS = weakly supported; 75–84% BS = moderately supported; and 85–100% BS = strongly supported; PP values ≥ 0.95: good support; PP values < 0.95: no support). Potential incongruence between datasets was assessed by visually inspecting the ML trees of individual gene regions. Well-supported incongruence was determined by strongly supported conflict with bootstrap values ≥ 85%. Incongruence of lower level of support was detected in some part of the topology. However, since no well-supported incongruence was detected in the backbone of the phylogeny among the nDNA trees and between cpDNA and nDNA datasets, the nDNA and cpDNA regions were concatenated for the above analyses.

### 2.4 Divergence time estimation

Fossil calibrations were implemented following recent recommendations and publications (Ho and Phillips, 2009; Pirie and Doyle, 2012; Massoni et al., 2015; Thomas et al., 2015; Chen et al., 2019). The Magnoliineae crown node was calibrated with a lognormal distribution (mean: 10.6, log(SD): 0.6; offset: 112.6,; median: ca. 121 Ma, 95% probability interval: ca. 136–112 Ma) with the hard minimum bound of ca. 112 Ma based on the age of *Endressinia brasiliana* Mohr & Bernardes-de-Oliveira (Mohr and Bernardes-de-Oliveira, 2004) and the soft upper bound based on the earliest angiosperm crown group pollen fossils from the Hauterivian (136.4–130 Ma; Friis et al., 2010). The Annonaceae crown node was calibrated with a lognormal distribution using *Futabanthus asamigawaensis* Takahashi et al., a Late Cretaceous (ca. 89 Ma) fossilized flower from Japan (Takahashi et al., 2008) to assign the hard minimum bound and *E. brasiliana* as the soft upper bound (mean 10.7; log(SD):0.6; offset: 89; median: ca. 98 Ma, 95% probability interval: ca. 112– 89 Ma).

Bayesian molecular divergence time estimation was performed in BEAST v.2.6.3 (Bouckaert et al., 2014) on the platform provided by the CIPRES Science Gateway (Miller et al., 2010). Conspecific duplicates were trimmed. Thirteen regions were partitioned based on the optimal partition scheme inferred by PartitionFinder v.2 (Lanfear et al., 2017). Yule process was specified as the tree prior (birth-death model failed to converge) in an uncorrelated lognormal relaxed molecular clock model (Drummond et al., 2006). A monophyly constraint was set to impose a sister relationship between Myristicaceae (*Coelocaryon preussii*) and Magnoliineae to avoid erroneous root topology (Pirie and Doyle, 2012). Four runs of MCMC analyses were performed, each with 140 million generations sampled every 10,000 generations.

Convergence of the MCMC output was assessed in Tracer v.1.7.1 (Rambaut et al., 2018) to ensure sufficient effective sample size (ESS > 200) for all parameters. The BEAST output trees were combined in LogCombiner v.2.4.6 (Bouckaert et al., 2014), with the first 25% of the samples of each run discarded as burn-in. The post-burn-in BEAST tree samples were summarized in maximum clade credibility (MCC) tree using in TreeAnnotator v.2.4.6 (Bouckaert et al., 2014).

### 2.5 Classification of fruit traits

Two fruit traits—fruit colour and monocarp width—were treated as binary characters. Fruit colour was coded as either ‘avian’ (A) or ‘non-avian’ (N), with avian colours defined according to the range of fruit colours preferred by birds but not limited only to birds (summarised by Voigt et al., 2004), including black, red, orange and yellow, whereas non-avian colours are restricted to brown and green. Monocarp width was coded as ‘narrow’ (N) or ‘wide’ (W). Beak gape size constrains most birds to the consumption of fruits that are narrower than 2.2 cm. Since a range of fruit maturity is present in herbarium specimens, and since drying often reduces monocarp width, ‘narrow’ is adopted for maximum monocarp widths < 2 cm, and ‘wide’ for those ≥ 2 cm. The fruit traits were assessed from taxonomic descriptions, photos, field observations and direct measurements from collections held in A, L, K, P, BISH, BKF, HKU, NOU and SING herbaria. Mature fruits were not available for three species due to the rarity of herbarium fruiting materials; in these cases, the character states were coded as ambiguous.

### 2.6 Trait-dependent historical biogeography

Maximum-likelihood ancestral range reconstruction was performed using BioGeoBEARS (Matzke, 2013; Klaus and Matzke, 2020). Ten areas were delimited based on the pattern of endemicity among extant *Meiogyne* species and palaeogeographical connectivity, viz.: (A) India; (B) SE Asia; (C) New Guinea; (D) Australia; (E) New Caledonia; (F) Fiji; (G) Tonga; (H) Vanuatu; (I) Mariana Islands; and (J) Neotropics. The maximum number of areas was set to three. Polyphyletic lineages of *M. cylindrocarpa* and *M. virgata* were treated as distinct species. Conspecific duplicates and outgroup taxa other than *Wuodendron* and the subtribe Sapranthinae were trimmed from the MCC tree. The accession *Meiogyne punctulata* 2 representing the synonym *M. tiebaghiensis* was trimmed due to a moderately supported conflict between cDNA and nDNA topologies.

BioGeoBEARS allows model comparison between DEC, DIVALIKE and BAYAREALIKE models by AICc weights across models. BioGeoBEARS also allows model variants with an extra free parameter +*j* to account for jump dispersal. Although the validity of the parameter *j* is controversial (Ree and Sanmartín, 2018; Matzke, 2022), we adopted +*j* models in the current study. To test whether fruit traits affect macroevolutionary dispersal of *Meiogyne*, trait-dependent models (*+t_12_ +t_21_ +m_2_*) and nested trait-independent models (*+t_12_ +t_21_*) were compared under the DEC, DIVALIKE and BAYAREALIKE models, where *t_12_* and *t_21_* are the character transition rates from binary state 1 to state 2, and vice versa (Klaus and Matzke). The parameter *m_1_*—the anagenetic dispersal rate multiplier at state 1—is a fixed parameter at the value 1 for all models, whereas *m_2_* describes the anagenetic dispersal rate multiplier at state 2 and was allowed to vary between 10 and 0. The *m_2_* value of 1 suggests the differential influence of trait states are negligible, while value < 1 signifies state 2 is associated with reduced dispersal rate, and conversely value > 1 suggests state 2 is correlated with elevated dispersal rate. Avian colour was coded as state 1 and non-avian colour as state 2, with narrow monocarps coded as state 1, and wide monocarps as state 2. The stepwise likelihood optimisation protocol suggested by Klaus and Matzke (2020) was adopted. A separate run was also performed in which the parameter *m_2_* was allowed to vary between 2 and 0 to facilitate likelihood optimisation. Statistical comparison was made across all models using AICc weights (Burnham and Anderson, 2004), coupled with likelihood ratio tests (LRTs) for comparisons among nested model pairs.

### 2.7 Ancestral character state reconstruction

One thousand ultrametric trees were resampled from the BEAST post-burn-in trees and were subsequently used as the input trees in Mesquite v.1.0 (Maddison and Maddison, 2003). Ancestral states were reconstructed using “Trace Character Over Trees” with AsymmMk models and were summarised on the MCC tree with the “Count Trees with Uniquely Best States” function.

### 2.8 Discrete character correlation

Correlation between fruit colour and monocarp width was tested using BayesTraits v.3.0 (Pagel and Meade, 2006). One thousand BEAST post-burn-in trees were used as the input trees for trait discrete dependent and independent models. All priors were set to an exponential with a mean of 10. To estimate the marginal likelihood, stepping stone analyses were implemented with 100 stones of 1,000 iterations each. Statistical comparison between the two models were made with log bayes factor which were interpreted as follows: < 2: weak evidence for H_1_; 2–5: positive evidence for H_1_; 5–10: strong evidence for H_1_; and > 10: very strong evidence for H_1_.

## 3 RESULTS

### 3.1 Phylogenetic analyses

The concatenated alignment of 60 accessions consists of 19,071 bp, of which 22.1% (4,211 bp) were variable and 7.8% (1,481 bp) were parsimony-informative (Appendix S4). The nDNA dataset has higher percentage of phylogenetically informative characters (PICs; 4.3%) than the cpDNA dataset (2.0%) within the ingroup. Within the ingroup, *NIA* possesses the highest number of PICs (69 bp), whereas *STG1* has the highest percentage of PICs (6.2%).

The cpDNA dataset yielded 10 equally most parsimonious trees of 1895 steps (consistency index, CI = 0.7689; retention index, RI = 0.6586; rescaled retention index, RC = 0.5064). The nDNA dataset generated 65 equally most parsimonious trees of 4162 steps (CI = 0.8044; RI = 0.7866; RC = 0.6328). The combined dataset yielded four equally most parsimonious trees of 6,118 steps (CI = 0.7854; RI = 0.7424; RC = 0.5831).

The topologies of the MP, ML, and Bayesian trees were largely similar within each dataset, with minor differences in the level of support (Fig. 1). The nDNA phylogeny confers much better resolution and robustness than the cpDNA phylogeny (Appendix S5). In the concatenated trees (Fig. 1), *Meiogyne* was retrieved as polyphyletic, with *Meiogyne papuana* nested within the well-supported genus *Monoon* (BS_MP_ = 100; BS_ML_ = 100; PP = 1). The remaining *Meiogyne* species were retrieved as a strongly supported clade (BS_MP_ = 100; BS_ML_ = 100; PP = 1), with four major clades identified. Clade A is the second most species-rich (BS_MP_ = 84; BS_ML_ = 88; PP = 1), comprising a larger Indomalayan subclade (A1: BS_MP_ = 99; BS_ML_ = 100; PP = 1) and a smaller Indochinese subclade (A2: BS_MP_ = 100; BS_ML_ = 100; PP = 1). Clade B is smaller, comprising three Indomalayan species ((BS_MP_ = 72; BS_ML_ = 58; PP = 1). A single species, *Meiogyne monosperma* (Hook. f. & Thomson) Heusden, forms Clade C. All the Australasian-Pacific species form Clade D (= APC), which is the most species-rich clade in *Meiogyne* (BS_MP_ = 100; BS_ML_ = 100; PP = 1). Within Clade D, the early divergent *M. cylindrocarpa* (D1) and four other strongly supported subclades were retrieved, including two Australian subclades (D3 and D4: BS_MP_ = 100; BS_ML_ = 100; PP = 1) and two New Guinea/Bismarck-Pacific subclades (D2: BS_MP_ = 83; BS_ML_ = 92; PP = 1; and D5: BS_MP_ = 100; BS_ML_ = 100; PP = 1, respectively), greatly improving the resolution of interspecific relationships within the APC. Clades C and D form a well-supported monophyletic group (BS_MP_ = 97; BS_ML_ = 99; PP = 1), although its relationships with the other two Indomalayan clades remain unresolved.

**Fig. 1.**
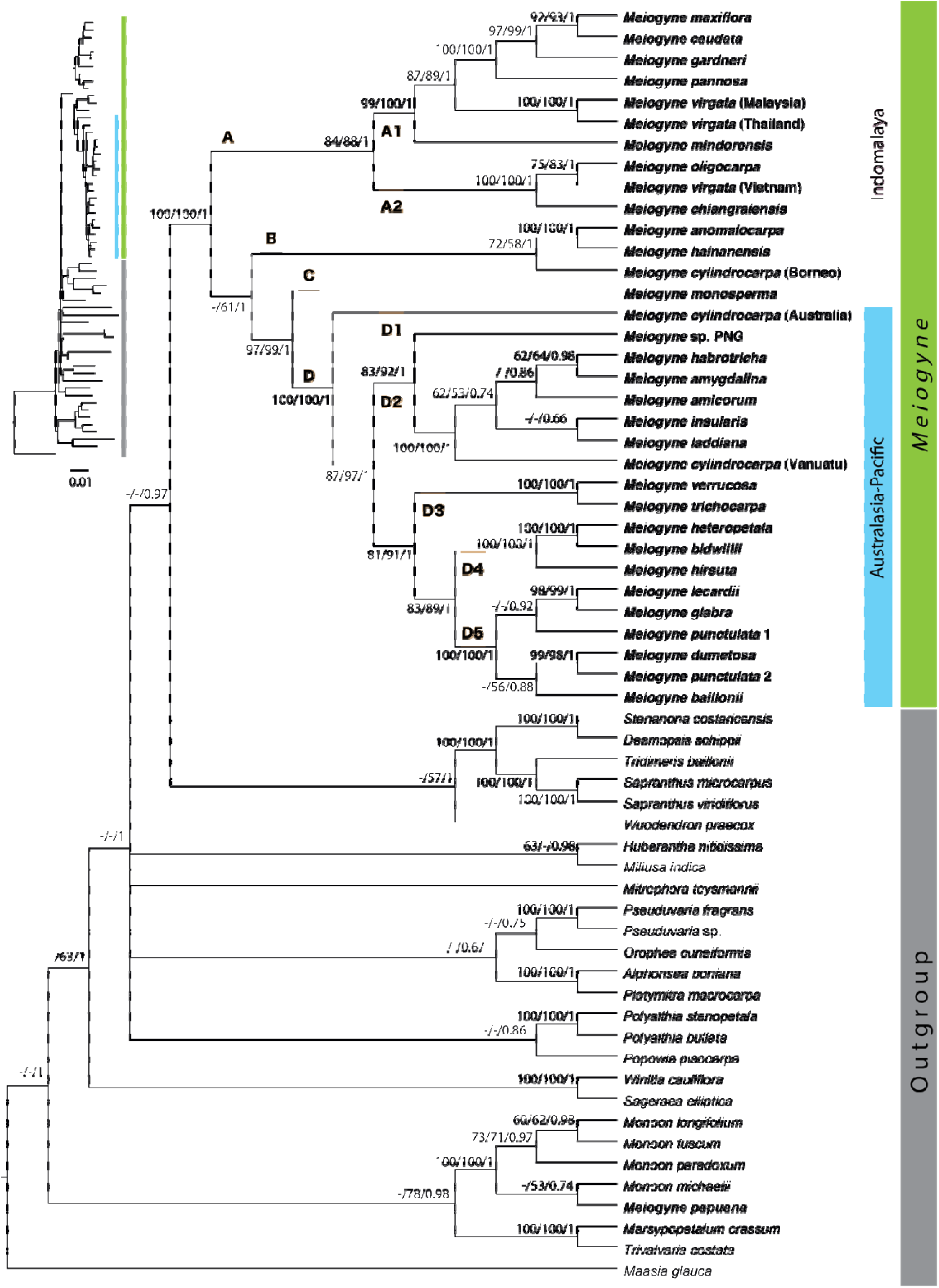
Bayesian 50% majority-rule consensus tree based on the complete concatenated datasets. The same consensus tree with branch lengths untransformed is shown on the left. Maximum parsimony bootstrap values (BS_MP_), maximum likelihood bootstrap values (BS_MP_) and Bayesian posterior probabilities (PP) were denoted at internal nodes in that order; − is annotated when BS_MP_, BS_ML_ or PP values are <50%.

The two widespread species, *M. virgata* and *M. cylindrocarpa*, are polyphyletic. The Vietnamese accession of *M. virgata*, (morphologically congruent with the type of its heterotypic synonym, *M. monogyna*), was nested within the well-supported Clade A2 (BSMP = 100; BSML = 100; PP = 1) instead of clustering with the other two *M. virgata* accessions. The Vanuatuan accession of *M. cylindrocarpa*, (cf. the type of its heterotypic synonym, *Oncodostigma wilsonii*), forms a strongly supported monophyletic group with all Fijian and Tongan species (BSMP = 100; BSML = 100; PP = 1). The Bornean accession of *M. cylindrocarpa* forms a moderately supported clade with two Indomalayan species (Clade B: BSMP = 72; BSML = 58; PP = 1). The New Caledonian species *M. punctulata* (Baill.) I.M.Turner & Utteridge is polyphyletic, with the accession similar to the type of *Meiogyne tiebaghiensis* (Däniker) Heusden (*Meiogyne punctulata* 2) retrieved as the sister of *Meiogyne dumetosa* (Vieill. ex Guillaumin) Heusden instead (BSMP = 98; BSML = 99; PP = 1).

### 3.2 Statistical comparison of biogeographic models

The BEAST MCC chronogram is shown in Appendix S6, and divergence time estimates and posterior probabilities of major internal nodes are summarised in Appendix S7. The parameter estimates, log likelihoods and AICc weights of all biogeographic models for fruit colour and monocarp width in BioGeoBEARS are summarised in Tables 1 and 2, respectively. For both trait datasets, trait-dependent models conferred the highest likelihood and AICc weight to the data. The best-fitting models for combined monocarp width and biogeographic data and combined fruit colour and biogeographic data are both the DEC +*j* +*t_12_* +*t_21_* +*m_2_* models (monocarp width: AICc: 161.50; AICc weight: 63.12%; fruit colour: AICc: 163.7; AICc weight: 45.30 %). For both traits, the best model performed significantly better than its trait-independent counterpart (monocarp width: LRT: *p* = 0.0016**; fruit colour: LRT: *p* = 0.044*). The anagenetic multiplier *m_2_* was 0.063 (fruit colour) and 0.049 (fruit width), suggesting fruits with narrow monocarp and avian colour were correlated with elevated macroevolutionary dispersal probability.

**Table 1.** Statistical comparison of biogeographic models incorporating fruit colour of *Meiogyne*. For LRT, *** denotes *p* < 0.0001; ** denotes *p* < 0.001 and; * denotes *p* < 0.05.

|  | *Geo*<br>*LnL* | *trait*<br>*LnL* | *LnL* | *d* | *e* | *j* | *t*<sub>12</sub> | *t*<sub>21</sub> | *m*<sub>2</sub> | *AICc* | *AICc*<br>Weight (%) | *p-value*<br>(adding *m*<sub>2</sub>) | *p-value*<br>(adding *j*) |
| --- | --- | --- | --- | --- | --- | --- | --- | --- | --- | --- | --- | --- | --- |
| *DEC* + *t*<sub>12</sub> + *t*<sub>21</sub> | -70.12 | -22.71 | -92.83 | 0.0027 | 0.001 |  | 0.059 | 0.054 | 1 | 194.9 | 0.00 |  |  |
| *DEC* + *j* + *t*<sub>12</sub> + *t*<sub>21</sub> | -53.74 | -22.71 | -76.46 | 0.0006 | 0.00 | 0.018 | 0.059 | 0.054 | 1 | 164.8 | 26.14 |  | 1.00E-08*** |
| *DIVALIKE* + *t*<sub>12</sub> + *t*<sub>21</sub> | -67.26 | -22.71 | -89.98 | 0.0031 | 0.0001 |  | 0.059 | 0.054 | 1 | 189.2 | 0.00 |  |  |
| *DIVALIKE* + *j* + *t*<sub>12</sub> + *t*<sub>21</sub> | -54.81 | -22.71 | -77.52 | 0.0007 | 0.00 | 0.018 | 0.059 | 0.054 | 1 | 167 | 8.70 |  | 6.00E-07*** |
| *BAYAREALIKE* + *t*<sub>12</sub> + *t*<sub>21</sub> | -104.72 | -22.71 | -109.6 | 0.003 | 0.037 |  | 0.059 | 0.054 | 1 | 228.5 | 0.00 |  |  |
| *BAYAREALIKE* + *j* + *t*<sub>12</sub> + *t*<sub>21</sub> | -103.01 | -22.71 | -78.33 | 0.0006 | 0.00 | 0.019 | 0.059 | 0.054 | 1 | 168.6 | 3.91 |  | 2.50E-15*** |
| *DEC* + *t*<sub>12</sub> + *t*<sub>21</sub> + *m*<sub>2</sub> |  |  | -89.33 | 0.0051 | 0.0033 |  | 0.061 | 0.048 | 0.00 | 190.6 | 0.00 | 0.0081** |  |
| ***DEC* + *j* + *t*<sub>12</sub> + *t*<sub>21</sub> + *m*<sub>2</sub>** |  |  | **-74.43** | **0.0011** | **0.00** | **0.029** | **0.059** | **0.052** | **0.063** | **163.7** | **45.30** | **0.044\*** | **4.80E-08\*\*\*** |
| *DIVALIKE* + *t*<sub>12</sub> + *t*<sub>21</sub> + *m*<sub>2</sub> |  |  | -86.74 | 0.0052 | 0.00 |  | 0.065 | 0.045 | 0.00 | 185.4 | 0.00 | 0.011* |  |
| *DIVALIKE* + *j* + *t*<sub>12</sub> + *t*<sub>21</sub> + *m*<sub>2</sub> |  |  | -75.85 | 0.0012 | 0.00 | 0.028 | 0.059 | 0.052 | 0.15 | 166.5 | 11.17 | 0.067 | 3.00E-06*** |
| *BAYAREALIKE* + *t*<sub>12</sub> + *t*<sub>21</sub> + *m*<sub>2</sub> |  |  | -105.4 | 0.0064 | 0.034 |  | 0.049 | 0.044 | 0.00 | 222.7 | 0.00 | 0.0035** |  |
| *BAYAREALIKE* + *j* + *t*<sub>12</sub> + *t*<sub>21</sub> + *m*<sub>2</sub> |  |  | -76.69 | 0.0009 | 0.00 | 0.03 | 0.059 | 0.052 | 0.15 | 168.2 | 4.78 | 0.07 | 3.60E-14*** |

**Table 2.**
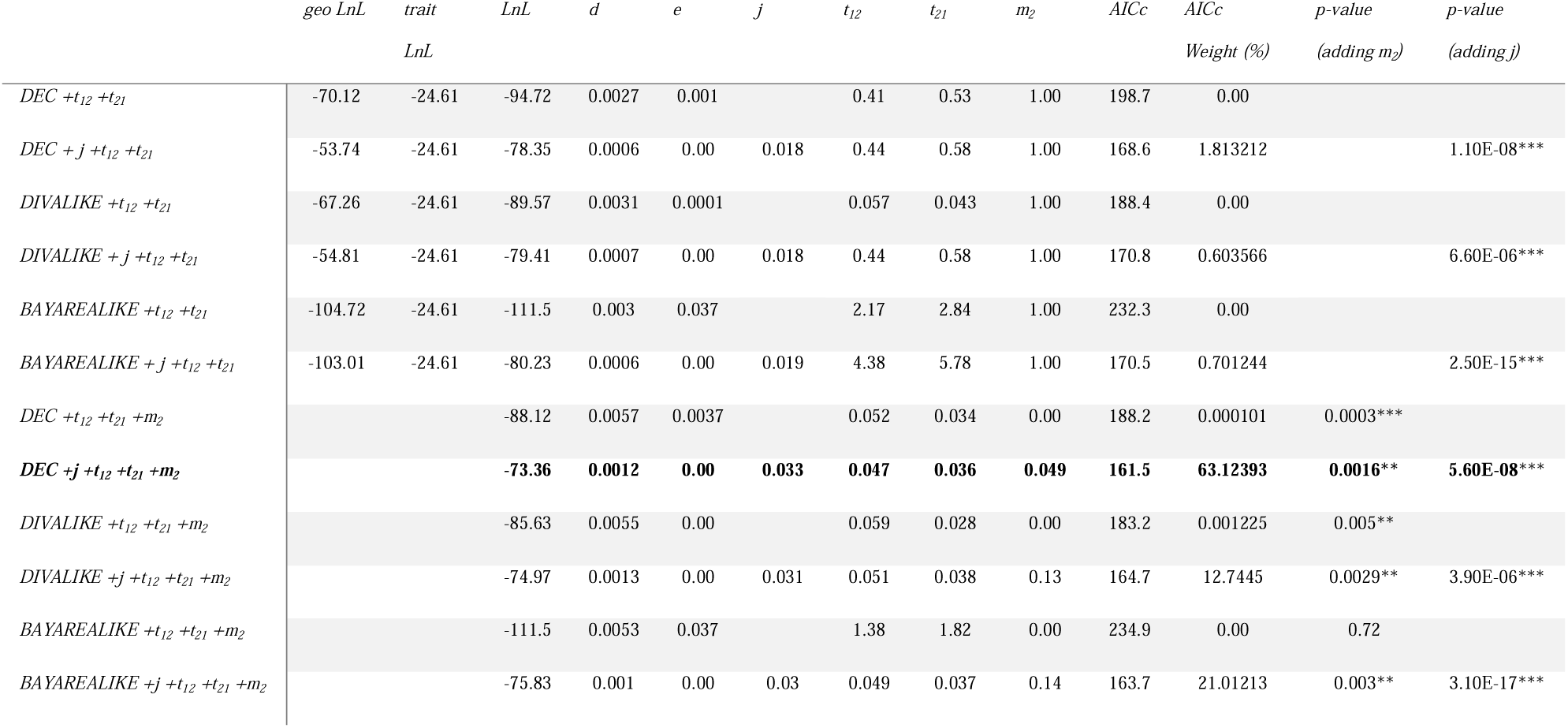
Statistical comparison of biogeographic models incorporating monocarp width of *Meiogyne*. For LRT, *** denotes *p* < 0.0001; ** denotes *p* < 0.001 and;* denotes *p* < 0.05.

For both traits, comparisons among models without the parameter +*j* retrieved the trait-dependent models DIVALIKE +*t_12_*+*t_21_* +*m_2_* as the best models (monocarp width: AICc weight: 86.4%; LRT: *p* = 0.005**; fruit colour: AICc weight: 81.1%; LRT: *p* = 0.011*), suggesting trait-dependent models conferred the highest likelihoods regardless of the parameter +*j*.

### 3.3 Ancestral range reconstruction

The ancestral range reconstruction is shown in Fig. 2. The best-fitting DEC +*j* +*t_12_* +*t_21_* +*m_2_* models for both traits yielded almost identical ancestral range inferences across the phylogeny. For simplicity, only the biogeographical model incorporating monocarp width is shown.

**Fig. 2.**
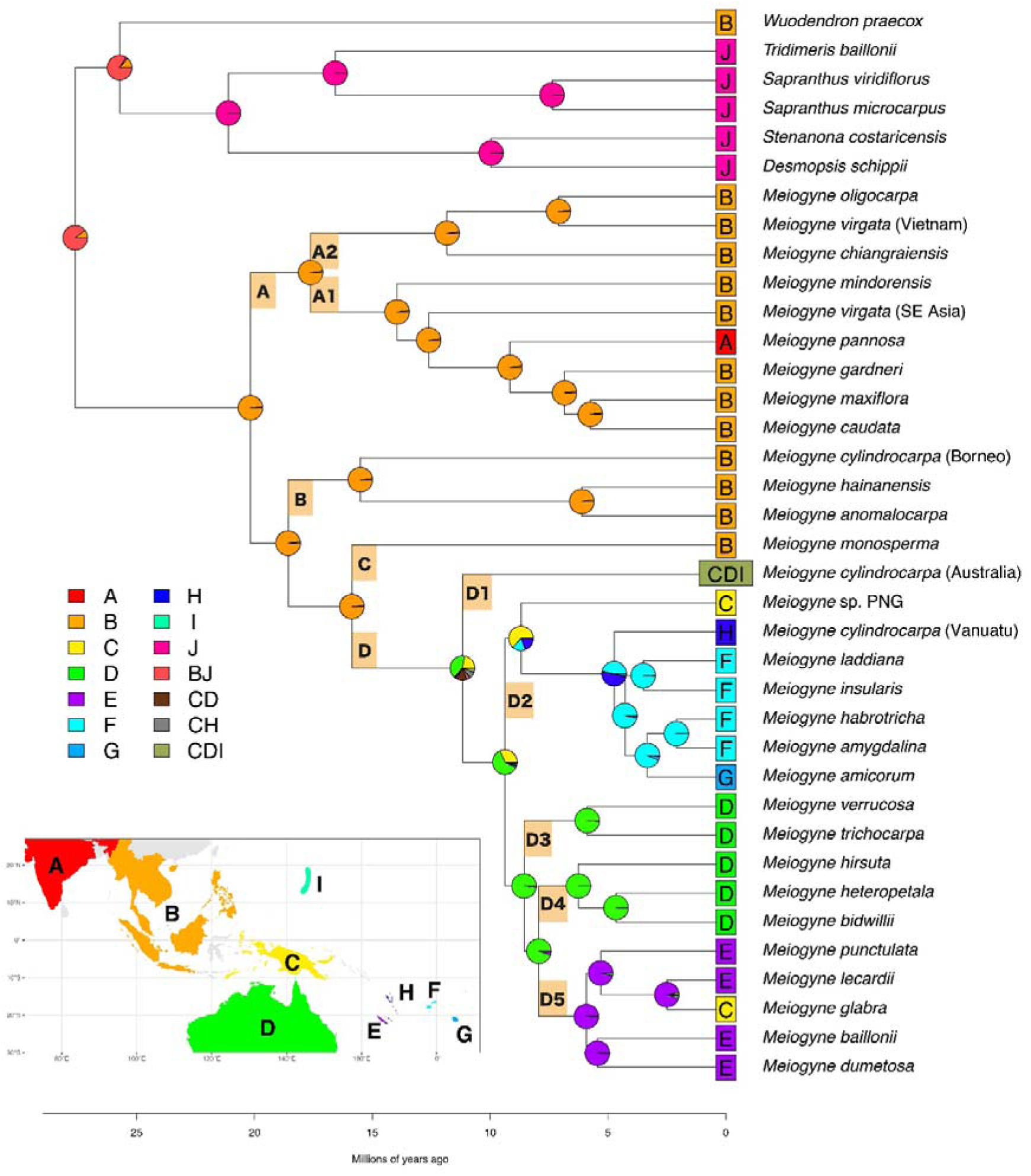
Trait-dependent historical biogeography of *Meiogyne* incorporating monocarp width under the best-fitting model (DEC +*j* +*t_12_* +*t_21_* +*m_2_*). Asterisk denotes the crown node for the Australasian-Pacific Clade (APC). Geographical area codes: (A) India; (B) SE Asia; (C) New Guinea; (D) Australia; (E) New Caledonia; (F) Fiji; (G) Tonga; (H) Vanuatu; (I) Mariana Islands; and (J) Neoetropics (not shown in the map).

The crown node of *Meiogyne* was shown to have occupied SE Asia (B). Most of the range evolution events occurred in the Australasia-Pacific Clade (APC). The ancestral range of the APC crown node, however, was equivocal and inferred to be New Guinea (C) or Australia (D) or the combination of both (C+D). Similarly, ancestral range estimation also yielded ambiguous results (either New Guinea and Australia) for the node immediately following the divergence of *M. cylindrocarpa*, suggesting the importance of Sahul (New Guinea + Australia) for subsequent dispersal into the Pacific islands.

Two Pacific clades were identified within the APC: the New Guinea-Pacific clade (Clade D2) and the New Guinea-New Caledonia clade (Clade D5). The former spans a wide range across the Western Pacific, including Vanuatu, Fiji and Tonga. New Guinea was shown to be the most probable ancestral range for the most recent common ancestor (MRCA) of the taxa in Clade D2, suggesting that New Guinea might be important for dispersal to the Pacific. On the other hand, New Caledonia was retrieved as the ancestral range of the MRCA of the Clade D5, where dispersal from Australia was responsible for colonisation of New Caledonia, suggesting Australia as another origin for dispersal from Sahul to the Pacific.

### 3.4 Ancestral character state reconstruction and character correlation

The ancestral states of the fruit colour and monocarp width traits were equivocal for the *Meiogyne* crown node and for most of the internal nodes in the Indomalayan grade (Fig. 3). Both traits were more evolutionarily labile in the Indomalayan grade. The MRCA of the APC and most of its internal nodes were estimated to have possessed fruits with narrow monocarps and fruit colour associated with bird dispersal. Bayesian character correlation models show strong correlation between the two monocarp traits (log marginal likelihood: independent: - 46.45; dependent: -41.95; log BF: 9.02).

**Fig. 3.**
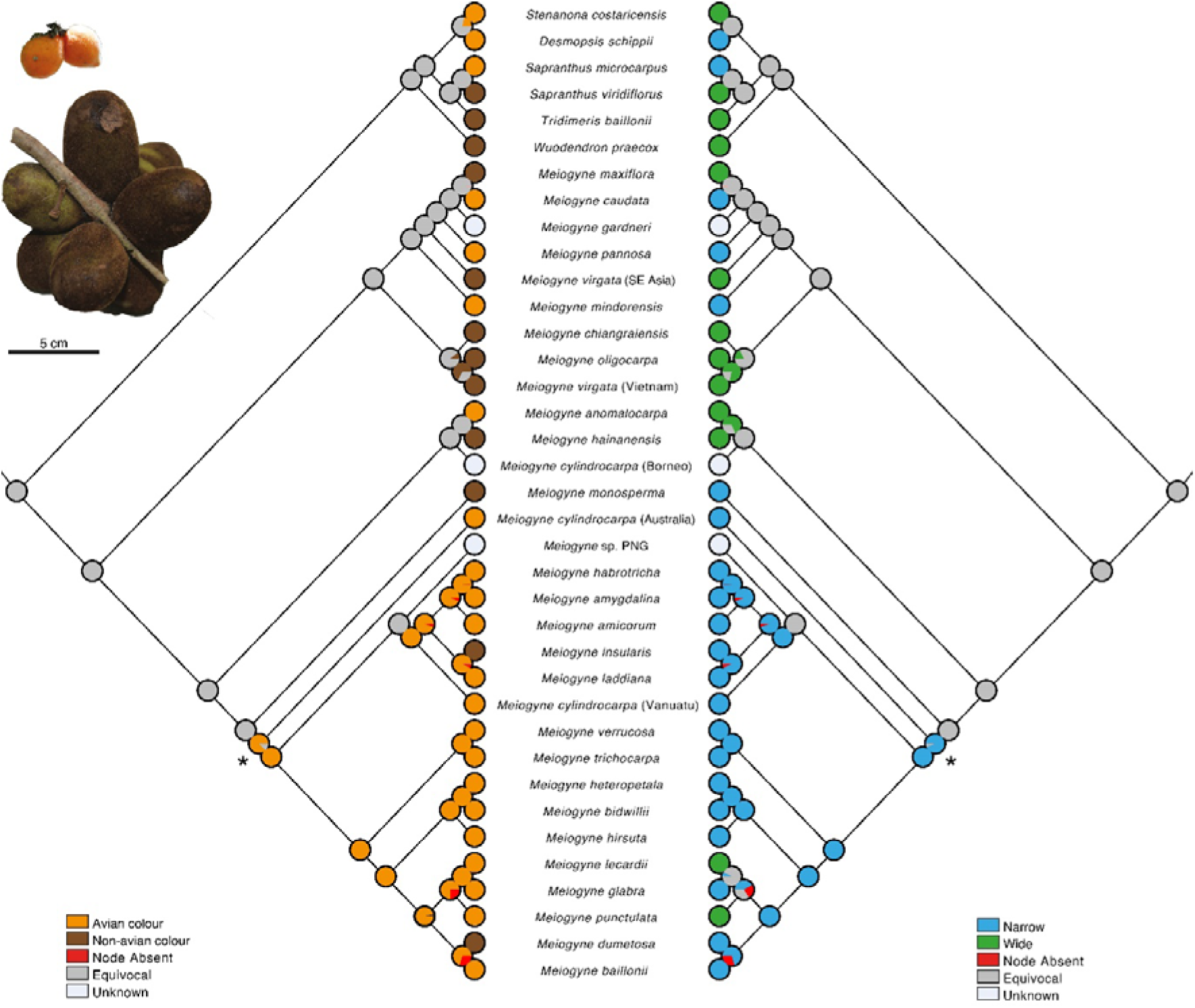
Ancestral character reconstruction of fruit colour (left) and monocarp width (right) in *Meiogyne*. Asterisk denotes the crown node for the Australasian-Pacific Clade (APC). Fruit from above to below: *Meiogyne heteropetala* (F.Muell.) D.C.Thomas, Chaowasku & R.M.K.Saunders and *Meiogyne maxiflora* D.M.Johnson & Chalermglin.

## 4 DISCUSSION

### 4.1 Phylogeny of *Meiogyne*

The combined nuclear and plastid phylogeny corroborates the known topology of *Meiogyne*, comprising a basal Indomalayan grade and a derived Australasian-Pacific Clade (Thomas et al., 2012; Xue et al., 2014), and demonstrates for the first time that the former comprises three clades. Morphological synapomorphies are not apparent for the major clades, precluding character diagnosis.

Molecular evidence here strongly suggests that *Meiogyne papuana* is nested within the well-supported genus *Monoon* (BS_MP_ = 100; BS_ML_ = 100; PP = 1), indicating that a taxonomic change is required to resolve paraphyly at the generic level. The species also possesses decurrent secondary vein insertion and a single seed per monocarp, which is characteristic of *Monoon* (Xue et al., 2012; Turner and Utteridge, 2015). *Meiogyne papuana* was previously placed in *Meiogyne* probably because of its inner petal corrugations, but rugosity at the base of the inner petals has been the source of taxonomic confusion due to its independent evolution in multiple lineages within the family, including *Asimina* Adans., *Alphonsea* Hook.f. & Thomson, *Duguetia* A.St.-Hil., *Orophea* Blume and *Pseuduvaria* Miq. (Xue et al., 2017b).

This study also provides molecular evidence for polyphyly in *M. cylindrocarpa*, *M. virgata* and *M. punctulata* at specific levels. Morphologically, the Vanuatuan specimens of *M. cylindrocarpa* are furthermore unique in their globoid to ovoid monocarps, while SE Asian specimens are distinguishable in habit, leaf shape and fruit size (Van Heusden, 1994). The northern Vietnamese accessions of *M. virgata* are distinguishable by reference to their adaxially glabrous petals and beaked, stipitate monocarps with constrictions between seeds. Finally, the New Caledonian *M. punctulata* shows great variation in leaf shape and size, as well as smooth or verruscose fruits, and probably represents yet another species complex. Polyphyly in these lineages could be more extensive and a taxonomic revision of the genus based on the examination of specimens across its range is required to address the problems arising from the polyphyly identified here.

### 4.2 Historical biogeography of *Meiogyne*

*Meiogyne* is one of the few genera in the Annonaceae that is particularly diverse in New Caledonia and the neighbouring Western Pacific Islands (Turner and Utteridge, 2017). This unusual distribution pattern has reinforced the hypothesis of a Gondwanan origin of the genus (Jessup, 1990; Lowry, 1998; Jaffré et al., 2001). This study unequivocally supports an alternative early-Miocene Indomalayan origin, however, as inferred in previous molecular phylogenetic studies of *Meiogyne* (Thomas et al., 2012; Xue et al., 2014). LDD of *Meiogyne* across Wallace’s line from West Malesia to Australasia occurred around the middle Miocene or the middle-late Miocene boundary. This was likely facilitated by the Sunda and Sahul collision around the Oligocene-Miocene boundary (Hall, 2001, 2002, 2009; Lohman et al., 2011). The resultant land mass, Wallacea, may have served as a ‘hopping platform’ for floristic dispersal across the strait, which might otherwise have been too difficult to cross. On the other hand, middle Miocene climatic cooling caused a fall in the global sea level, and the emergence of the continental shelf and previously submerged islands likely further facilitated macroevolutionary dispersal across Wallace’s line (Flower and Kennett, 1994; Scotese, 2014).

Ancestral range reconstructions suggest that Sahul was likely a stepping stone for subsequent macroevolutionary dispersal of *Meiogyne* into the Western Pacific. The two distinct Pacific clades likely resulted from two independent dispersal events from Sahul to the Pacific. Colonisation of the New Guinea-Pacific Clade D2 probably occurred from New Guinea to Vanuatu, Fiji and Tonga with island-hopping along the Vitiaz arc system (Crawford et al., 2003; Schellart et al., 2006). This hypothesis however requires an additional local extinction to explain the disjunction at the Solomon Islands. In contrast, dispersal to New Caledonia between the stem and crown node of Clade D5, dated around the middle to late Miocene, likely have originated from Australia instead of New Guinea. Similar dispersal pattern from Australia to New Caledonia is commonly documented in the literature, such as in Sapotaceae (Bartish et al., 2011) and Sapindaceae (Buerki et al., 2012). The occurrence of a single Clade D5 species in New Guinea, however, appears to be due to secondary dispersal from New Caledonia. It is interesting to note that reports on dispersal from New Caledonia to other islands is considerably rarer than dispersal into the archipelago, and consists of mostly dispersal towards Vanuatu (Pillon, 2011; Plunkett and Lowry, 2012; Munzinger et al., 2022).

Ancient global sea level changes might have assisted the dispersal into the Pacific. The fall in the global sea level associated with the middle Miocene cooling is likely to have lasted until the end of the Tortonian (ca. 7.246 Mya; Flower and Kennett, 1994; Scotese, 2014). This was succeeded by a global sea-level decline caused by the Messinian salinity crisis until the end of the late Miocene, with the sea level in South Pacific decreasing by at least 30 m (5.26 Mya: Aharon et al., 1993; Krijgsman et al., 1999). These two events coincided with the window of dispersal to New Caledonia and likely have exposed the Chesterfields and the associated Bellona Plateau and Kenn Plateau located between Australia and New Caledonia (Scotese, 2014), providing a stepping stone for island hopping. The sea-level fall may have further enabled LDD by exposing the Queensland and Marion Plateau off the Australian coast. A similar colonisation route has been suggested for peacock swallowtail butterflies (Condamine et al., 2013).

Many New Caledonian species have evolved an edaphic preference for ultramafic soil owing to the abundance of peridotite and serpentinite minerals on the archipelago. Interestingly, the New Caledonian Clade D5 is sister to the Australian Clade D4, which occurs along the eastern coast of Australia, adjacent to serpentine formations in the Marlborough terrane in central coastal Queensland (Murray, 1969, 2007; Thomas et al., 2012; Van der Ent et al., 2015). The ultramafic habitats there might have allowed pre-adaptation to ultramafic soil prior to transoceanic dispersal, potentially improving seedling recruitment and chances of population establishment. This hypothesis requires further scrutiny using ancestral niche reconstruction to assess whether evolution of ultramafic tolerance predated the dispersal event. Nonetheless, pre-adaptation and exaptation for ultramafic tolerance have been suggested to contribute significantly to the success of over-represented flora in ultramafic habitats (Pillon et al., 2010).

### 4.3 Fruit character correlation

Frugivorous birds exhibit the strongest preference for black and red fruits (Janson, 1983; Willson et al., 1990). They nevertheless consume some fruits with a broader range of colours, including those that are orange, yellow, purple, white, blue and pink (Knight and Siegfried, 1983; Voigt et al., 2004). Most bird-dispersed *Meiogyne* fruits are black, red and orange (Wandrag et al., 2015; Pohlman, 2006; Ganesh and Davidar, 2001). While black and red fruits provide honest antioxidant rewards (Schaefer et al., 2008), yellow and orange fruits are associated with higher protein levels and lower tannin content (Schaefer and Schmidt, 2004; Schaefer and Ruxton, 2011). Birds might consume yellow and orange fruits because they provide a more reliable source of macronutrients to fulfill their dietary requirements.

Irrespective of diffuse evolution, fleshy fruits often show a suite of morphological adaptions to frugivore guilds: for example, mammalian dispersal syndromes are often characterised by large and dull-coloured fruits with a protective husk or thick pericarp, whereas avian dispersal syndromes are characterised by small fruits with thin pericarp of specific colours, including red, black, orange, purple, yellow, white, pink and blue (Janson, 1983; Knight and Siegfried, 1983; Gautier-Hion et al., 1985; Voigt et al., 2004; Chen et al., 2020). It is not surprising that in *Meiogyne* narrow monocarps are correlated with these bright fruit colours since these functional characters are involved in seed dispersal and are associated with this dispersal syndrome (Voigt et al., 2004). Association between larger fruits and duller pigmentation could also be due to metabolic cost constraints (Schaeffer and Ruxton, 2011): higher metabolic costs associated with anthocyanin biosynthesis are inevitably incurred to produce the same level of pigmentation in larger fruits, although larger fruits are intrinsically salient.

### 4.4 Influence of fruit traits on macroevolutionary dispersal

Available data on frugivory in *Meiogyne* largely aligns with the dispersal syndrome hypothesis: species with dull-coloured and wide monocarps, such as *M. hainanensis* and *M. virgata*, are reportedly consumed by gibbons (Elder, 2013; Liu, 2015, as “*Oncodostigma*”), whereas birds are the only known dispersal agents for species with brightly coloured and narrow monocarps, such as *M. cylindrocarpa* (Wandrag et al., 2015), *M. heteropetala* (pers. obs.), *M. hirsuta* (Pohlman, 2006, as “*Meiogyne* sp. Henrietta Creek LWJ 512”) and *M. pannosa* (Ganesh and Davidar, 2001). Our BioGeoBEARS analysis for *Meiogyne* shows that the fruit traits related to bird dispersal—narrow monocarp width and avian fruit colour—are correlated with increased macroevolutionary dispersal, suggesting that avian dispersers might have promoted LDD across oceanic barriers. Unlike most mammals, the size of fruits consumed by birds are restricted by their gape size (Wheelwright, 1985). With only a few notable exceptions (fruit pigeons and hornbills), most birds are restricted to the consumption of fruits that are narrower than 2.2 cm (Leighton and Leighton, 1983). The almost exclusively west-to-east dispersal in *Meiogyne* is congruent with the migratory pattern of birds in the East Asian-Australasian flyway (Nathan et al., 2008; Gillespie et al., 2012), suggesting that migratory frugivorous birds may have been an important dispersal agent for assisting LDD in *Meiogyne*.

Nonetheless, the effect of fruit traits on LDD generally appears to be highly context-specific, especially in Annonaceae (Onstein et al., 2019) and are probably heavily influenced by palaeogeography, ancestral frugivore communities and phylogenetic lineages. Diaspore traits related to bird dispersal, such as red pigmentation, pericarp fleshiness and moniliform fruits, have been shown to promote anagenetic dispersal capability in Ericaceae tribe Gaultherieae (Lu et al., 2019), Podocarpaceae (Klaus and Matzke, 2020), and Annonaceae (Onstein et al., 2019). Although the effect of fruit width on macroevolutionary dispersal is yet be tested in the framework of trait-dependent historical biogeography, larger syncarpous fruits in Annonaceae, associated with megafauna and large mammal dispersal, have been shown to promote LDD in Laurasia (ca. 66–45 Ma) and across South America and Africa (ca. 66–33.9 Ma) (Onstein et al., 2019), but do not assist in Pacific Island hopping. This is congruent with our findings and can be attributed to our narrower scope in Indomalaya and the Australasia-Pacific region where megafauna and larger mammals rarely crossed Wallace’s line (Webb, 2013), thus limiting the importance of mammal-dispersal fruit traits in transoceanic seed dispersal.

Caution is nevertheless essential since trait-dependent biogeographical models do not detect causation but merely correlation between traits and macroevolutionary dispersal capability. Underlying causes for the traits may instead be responsible for the observed elevated anagenetic dispersal rates. The traits could also be products of adaptation to novel geographical area and niches: for example, fruit size has been shown to co-vary with leaf size (Herrera, 2002), which might have been an adaptation for island climate.

## 5 CONCLUSIONS

Questions regarding the effect of traits on biogeographical pattern have long been raised (Darwin, 1859). Recent advances in trait-dependent historical biogeography have laid a framework for statistical model comparisons to test the effects of traits in macroevolutionary dispersal, which were previously elusive (MatoslMaraví et al., 2018; Klaus and Matzke, 2020). Our findings suggest that avian dispersal-related character states—narrow monocarp width and avian fruit colour—promoted macroevolutionary dispersal in the tropical woody lineage *Meiogyne.* The traits tested (fruit width and colour) are universal and the findings can therefore easily be related to other endozoochorous plant lineages: this study therefore has the potential to shed light on the interactions between traits and biogeography in other lineages with similar dispersal patterns.

## ACKNOWLEDGEMENTS

Financial support was provided by grant HKU17112616 from the Hong Kong Research Grants Council, awarded to RMKS. We are grateful to Dr Charan Leeratiwong, Dr Michael Thomas, Dr Shao Yun Yun, and Dr Richard Chung Cheng Kong for providing leaf material, and Laura Wong for providing general technical support.

## DATA AVAILABILITY STATEMENT

All DNA sequence data used in this study are available from the nucleotide database of the National Center for Biotechnology Information (http://www.ncbi.nlm.nih.gov), with GenBank accession numbers and locality data listed in Appendix S8.

