## Supplemental Data 1 for "Trait-dependent biogeography offers insights on the dispersal of *Meiogyne* (Annonaceae) across the Australasia-Pacific region"

| Gene Region | Primer | Sequence (5’->3’) | Source |
| --- | --- | --- | --- |
| *ATPQ* (intron4) | ATPQ c393F | GAGATCCCCAAGTATGTAGACA | This study |
|  | ATPQ d311R | CAGCAATTTCCTTTTCCAGC | This study |
|  | ATPQ de527R | GTCTACCACATGCAAATATTG | This study |
|  | ATPQ d242F | TTGGTGGAATTGAAAGAAGC | This study |
|  | ATPQ de345R | ATTTCTGCTTCCCTGACCTT | This study |
|  | ATPQ de398F | CTGTCCTATCAGTGCTACCT | This study |
|  | ATPQ ce660R | GTGGACAACAGGATAGCAG | This study |
|  | ATPQ ce579F | GGAGGGATCAGGTTTATCTTG | This study |
|  | ATPQ e587R | AGGATGCTTCTCAAAATACTCA | This study |
| *COX6A* (intron 1) | COX6A a41F | GATAGCAAGGTTGGGAATTCA | This study |
|  | COX6A ab308R | ATCTCAAACAAACAGTGGCC | This study |
|  | COX6A ab184F | GGGGATTCTGYTGTGTTTTCTA | This study |
|  | COX6A ab487R | ATTTGCATGAATTGAACACTAGT | This study |
|  | COX6A ab371F | AACTGTCAAACGCAGGATG | This study |
|  | COX6A ab724R | CTTGGCATGTCTACTGCAG | This study |
|  | COX6A ab529F | GCCATGTTGCTTTGGTACT | This study |
|  | COX6A ab842R | AACAACACAAAGAAAAGGATAACC | This study |
|  | COX6A ab706F | CTGCAGTAGACATGCCAAG | This study |
|  | COX6A ab1009R | TAGGTGGTAAATTGACAGAACG | This study |
|  | COX6A ab909F | ATTTATTTGTTTGGTTTCTATGTTAGA | This study |
|  | COX6A b215R | TAGGTGGTAAATTGACAGAACG | This study |
|  | COX6A d325R | TCCTTGTGCTTCTTCTCAAAC | This study |
| *CHER1* (intron 4) | CHER1 d890F | ACAGTCTTCCTCTTTGATGTTC | This study |
|  | CHER1 de457R | CATTTTCCAACCTTGCAACC | This study |
|  | CHER1 de365F | GAATGCTTAAATACCTGGCAAG | This study |
|  | CHER1 de693R | ACATCCTACAAACTGCAAAGAG | This study |
|  | CHER1 de630F | CTAGAGCTTCCCTATTCCTTAT | This study |
|  | CHER1 de916R | ACATAGTCACTATTTCCTTTTTG | This study |
|  | CHER1 de839F | ATAGCTTGGGGATCCTTCAG | This study |
|  | CHER1 e974R | ATACAGCATCATGTCCAATCC | This study |
| *DDB2* (exons 1-3 and intervening introns) | DDB2 a259F | AATCACCATCAGCCTCAAGAAA | This study |
|  | DDB2 b377R | GCATTTTACAGAGAAAGCAAGG | This study |
|  | DDB2 b318F | GATTCAAGGGTGCTACTTACAT | This study |
|  | DDB2 c483R | CGCTCAAACACATAATCCAAG | This study |
|  | DDB2 abc486R | GGCATGTCATTGTGGTATGG | This study |
| *DJC65* (intron 2) | DJC65 a596F | CGTTTCAATCCAAAAAGAACGA | This study |
|  | DJC65 bc344R | GCATGCTTGGATAGTGATAAAA | This study |
|  | DJC65 bc266F | ACTCTATAGCTAACATCACCCT | This study |
|  | DJC65 bc526R | ACAGCTAATGAACCATCTTGAA | This study |
|  | DJC65 bc447F | GGTTTAATTTATGACATTGTTCAGC | This study |
|  | DJC65 b722R | CCAGCCTTCTATAAGCAGATTT | This study |

| Gene Region | Primer | Sequence (5’->3’) | Source |
| --- | --- | --- | --- |
| *EIF3K* (intron 3) | EIF3K c343F | CTCCAGATTTCAACCTTTGTCT | This study |
|  | EIF3K cd374R | AAAAGGGGCAAAAKGTGTG | This study |
|  | EIF3K cd318F | CCACTGTTGATGTTTTGTAGTC | This study |
|  | EIF3K cd411F | AGGCTGTTTGTTGAATGAATCT | This study |
|  | EIF3K cd621R | GCATGTTATTCTCTCCATGAAC | This study |
|  | EIF3K d436R | TCCAGGTAGTGTGAAAGCA | This study |
| *MNJ7.16* (exons 1-3 and intervening introns) | nduB8 a1F | GAAAATGGCGGGAAATTTGAG | This study |
|  | nduB8 b326R | GTGCGAGACGATCAATGC | This study |
|  | nduB8 b253F | GGTGCCTGTAAACGATGAG | This study |
|  | nduB8 c323R | AGGGAATCTTAGACGCTTTATC | This study |
| *NIA* (intron 2) | NIA-i2 F | TCBGTGATTACGACGCCGTGTCATGA | Howarth & Baum, 2002 |
|  | NIA-i3 R | GAACCARCARTTGTTCATCATDCC | Howarth & Baum, 2002 |
|  | NIA b1F | GATATGCATATTCCGGTGAGTG | This study |
|  | NIA bc336R | TAATTCTACCATCGGATCTACC | This study |
|  | NIA bc232F | GACGGTCTATATGGATGGAGA | This study |
|  | NIA bc510R | TTTTAGATTCCAAACCCGCTT | This study |
|  | NIA bc450F | TGAGAAGTCTGTCCAACGA | This study |
|  | NIA bc724R | CGTGCTACATTAACATACATGC | This study |
|  | NIA bc620F | GATAACTGCTACACACGCC | This study |
|  | NIA bc943R | AAAATGGGGATTAAAATGCAAT | This study |
|  | NIA bc810F | AGGCATAAAGTAAACAAAAATAGAT | This study |
|  | NIA bc1139R | ATGTTGTTTGTCGGATTCCAT | This study |
|  | NIA bc1049F | CTTTTTGTCAGGATCCATCATC | This study |
|  | NIA bc1365R | ATTTTAATATTCAACTGCGCCA | This study |
|  | NIA bc1278F | TGCACTTGTCCAAATATGTTCT | This study |
|  | NIA bc1547R | CAGAAACACCAGCAATAGTACT | This study |
| *PDF5* (intron 2) | PFD5 b376F | GAAACTGTCCTTGTTGATGTTG | This study |
|  | PFD5 bc333R | GCTTTTTACAATGGCTACTGTT | This study |
|  | PFD5 bc234F | AATTCAGTTCAATGTAGAGGYT | This study |
|  | PFD5 bc535R | CAAGTTACAACATKCCRCAC | This study |
|  | PFD5 bc457F | GCAAGTTAATAGGGAAACATTAGC | This study |
|  | PFD5 bc674R | ACCTTCCCCACTACATTATGAT | This study |
|  | PFD5 bc576F | TTGTGGTAATTATGGGGGAATG | This study |
|  | PFD5 bc815R | TAGTTTCCTTTTTCCTCCCACT | This study |
|  | PFD5 bc794F | AGTGGGAGGAAAAAGGAAAC | This study |
|  | PFD5 bc1118R | TCAGAAGAAAAGGAAGAAAATAGAA | This study |
|  | PFD5 bc1006F | ATTTGGGCCAGTTTTCTTTTAG | This study |
|  | PFD5 c481R | CGTAGTTTGACTTAAGGAGGTT | This study |

| Gene Region | Primer | Sequence (5’->3’) | Source |
| --- | --- | --- | --- |
| *STG1* (intron 5) | STG1 e419F | ATAAGCGCTTGATCTTGACAA | This study |
|  | STG1 ef308R | CAGAAATGACAAGTTATGCTACRA | This study |
|  | STG1 ef249F | TGGATCATGAAACACGTATACA | This study |
|  | STG1 f495R | TTCTTGGTGTTTCATATTCACG | This study |
| *MHF15.12* (exons 6-9 and intervening introns) | s8e38F | GTGGCTGAAGATGAAATGTT | This study |
|  | s8e M315R | GCCCTACAAATGATATAAGAGC | This study |
|  | s8e M208F | CAATCTGCACTATGGGCAAG | This study |
|  | s8e M514R | AGCTGGATCTAACTATTGTAAGAC | This study |
|  | s8e M408F | GAGTGAGGCATTCTTAAGTGTT | This study |
|  | s8e M692R | ATTTCAGCTCTGGATGGGT | This study |
|  | s8e M604F | CACTAGAAAACCTCCAAAATATGA | This study |
|  | s8e M841R | TGCTTGTAAGAAAACACTACCT | This study |
|  | s8e M738F | GTCCCATGTACACTTCTCTTG | This study |
|  | s8e M1067R | ATCTCATTCACAATAGGCCATC | This study |
|  | s8e M993F | TGGGGTAAGATTTGAGACATTC | This study |
|  | s8e M1258R | AGCACATTGGAGGATCAAAC | This study |
|  | s8e M1157F | GAATCATCAAGTGTTCCGAAG | This study |
|  | s8e480R | TTCTCTGGATTATTTGTGACTTG | This study |

**Appendix S1** List of primers used for amplification and sequencing of 11 low-copy nuclear genes. The numbering of the introns and exons was based on that of *Arabidopsis thaliana.*

| **Dataset** | **Partition** | **Model selected** | |
| --- | --- | --- | --- |
|  |  | **RAxML** | **MrBayes** |
| **cpDNA** | *matK*, ndhF | GTR + Γ | GTR + I + Γ |
|  | *ndhF-rpl32, rpl32-trnL* |  | GTR + I + Γ |
|  | *rbcL* |  | GTR + I + Γ |
|  | *trnLF* |  | HKY + Γ |
|  | *ycf1* |  | GTR + I + Γ |
| **nDNA** | *ATPQ, CHER1, STG1* | GTR + Γ | GTR + Γ |
|  | *COX6A, DDB2, DJC65, EIF3K* |  | GTR + Γ |
|  | *MNJ7.16* |  | HKY + Γ |
|  | *NIA* |  | GTR + Γ |
|  | *PFD5, MHF15.12* |  | HKY + Γ |
| **cpDNA + nDNA** | *matK*, ndhF | GTR + Γ | GTR + I + Γ |
|  | *ndhF-rpl32, rpl32-trnL* |  | GTR + I + Γ |
|  | *rbcL* |  | GTR + I + Γ |
|  | *trnLF* |  | HKY + Γ |
|  | *ycf1* |  | GTR + I + Γ |
|  | *ATPQ, STG1* |  | HKY + Γ |
|  | *COX6A, DDB2* |  | GTR + Γ |
|  | *CHER1, DJC65, EIF3K* |  | GTR + Γ |
|  | *MNJ7.16* |  | HKY + Γ |
|  | *NIA* |  | GTR + Γ |
|  | *PFD5, MHF15.12* |  | HKY + Γ |

**Appendix S2** Partitions and molecular substitution models used for the RAxML and MrBayes analyses.

| **Dataset** | **Partition** | **Model selected** |
| --- | --- | --- |
| **cpDNA + nDNA** | *matK* | GTR +G +X |
|  | *ndhf* | GTR +I +G +X |
|  | *ndhf-rpl32, rpl32-trnL* | GTR +G +X |
|  | *rbcL* | GTR +I +G +X |
|  | *trnLF* | GTR +G +X |
|  | *ycf1* | GTR +I +G +X |
|  | *ATPQ* | HKY +G +X |
|  | *COX6A , DDB2* | GTR +G +X |
|  | *CHER1, DJC65, EIF3K* | GTR +G +X |
|  | *MNJ7.16* | HKY +G +X |
|  | *NIA* | GTR +G +X |
|  | *PFD5, MHF15.12* | HKY +G +X |
|  | *STG1* | GTR +G +X |

**Appendix S3** Partitions and molecular substitution models used for the BEAST analyses.

| **DNA region** | **Alignment**  **Length (bp)** | **Missing data (%)** | | **Variable characters (%)** | | **Parsimony-informative characters (%)** | |
| --- | --- | --- | --- | --- | --- | --- | --- |
|  |  | **Entire**  **dataset** | **Ingroup** | **Entire**  **dataset** | **Ingroup** | **Entire**  **dataset** | **Ingroup** |
| **Chloroplast DNA data** | | | | | | | |
| *matK* | 770 | 0.8 | 1.5 | 139 (18.1) | 56 (7.3) | 50 (6.5) | 17 (2.2) |
| *ndhF* | 2037 | 15.1 | 9.7 | 298 (14.6) | 96 (4.7) | 114 (5.6) | 33 (1.6) |
| *ndhf-rpl32* | 667 | 30.1 | 16.6 | 144 (21.6) | 65 (9.7) | 50 (7.5) | 28 (4.2) |
| *rbcL* | 1378 | 14.6 | 19 | 105 (7.6) | 34 (2.5) | 36 (2.6) | 17 (1.2) |
| *rpl32-trnL* | 1328 | 27.6 | 6.5 | 235 (17.7) | 128 (9.6) | 68 (5.1) | 48 (3.6) |
| *trnL-F* | 912 | 4.9 | 5 | 141 (15.5) | 44 (4.8) | 43 (4.7) | 18 (2.0) |
| *ycf1* | 1598 | 8.4 | 10.8 | 263 (16.5) | 74 (4.6) | 55 (3.4) | 16 (1.0) |
| Combined data | 8690 | 14.6 | 9.4 | 1325 (15.2) | 497 (5.7) | 416 (4.8) | 177 (2.0) |
| **Nuclear DNA data** | | | | | | | |
| *ATPQ* | 622 | 12.3 | 11.8 | 295 (47.4) | 82 (13.2) | 119 (19.1) | 33 (5.3) |
| *COX6A (cco)* | 1235 | 58.0 | 36.2 | 208 (16.8) | 139 (11.3) | 62 (5.0) | 37 (3.0) |
| *CHER1* | 1033 | 42.6 | 18.1 | 262 (25.4) | 149 (14.4) | 94 (9.1) | 58 (5.6) |
| *DDB2* | 555 | 21.4 | 25.7 | 220 (39.6) | 60 (10.8) | 79 (14.2) | 17 (3.1) |
| *DJC65* | 732 | 13.9 | 15.6 | 294 (40.2) | 103 (14.1) | 127 (17.3) | 43 (5.9) |
| *EIF3K* | 675 | 22.6 | 17.1 | 252 (37.3) | 78 (11.6) | 89 (13.2) | 28 (4.1) |
| *MNJ7.16* | 545 | 12.8 | 11.0 | 194 (35.6) | 66 (12.1) | 81 (14.9) | 29 (5.3) |
| *NIA* | 1563 | 47.6 | 19.5 | 341 (21.8) | 242 (15.5) | 110 (7.0) | 69 (4.4) |
| *PDF5* | 1313 | 43.0 | 18.1 | 279 (21.2) | 170 (12.9) | 78 (5.9) | 40 (3.0) |
| *MHF15.12* | 1509 | 49.9 | 21.8 | 241 (16.0) | 187 (12.4) | 91 (6.0) | 60 (4.0) |
| *STG1* | 599 | 11.1 | 19.2 | 288 (48.1) | 93 (15.5) | 135 (22.5) | 37 (6.2) |
| Combined data | 10381 | 36.9 | 20.5 | 2886 (27.8) | 1369 (13.2) | 1065 (10.2) | 451 (4.3) |
| **Combined chloroplast and nuclear data** | | | | | | | |
| Combined data | 19071 | 40.6 | 15.7 | 4211 (22.1) | 1866 (9.8) | 1481 (7.8) | 628 (3.3) |

**Appendix S4** Descriptive statistics for the seven cpDNA and 11 nDNA markers and the concatenated datasets.

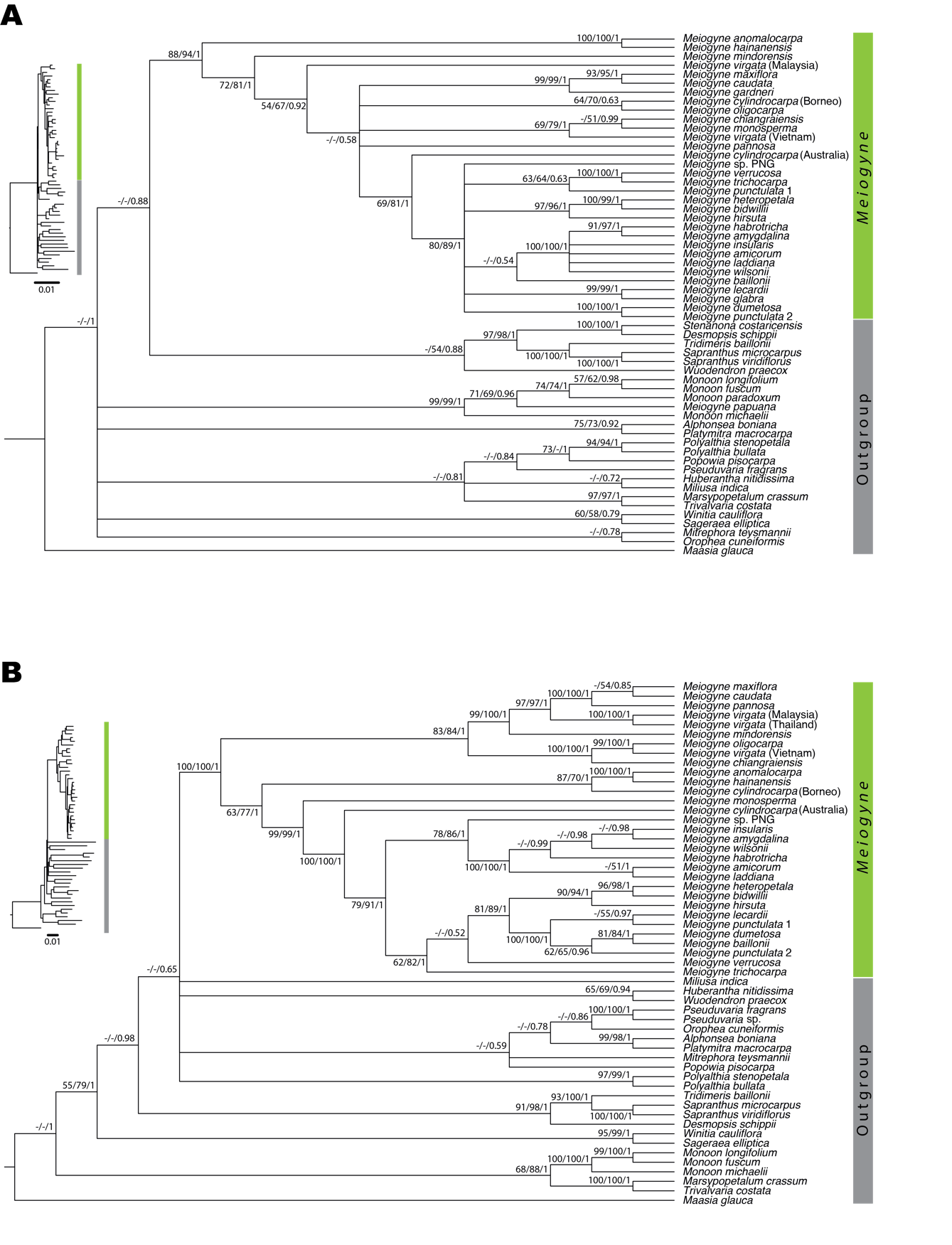

**Appendix S5** Bayesian 50% majority-rule consensus tree based on individual datasets. A) based on concatenated dataset of seven cpDNA regions. B) based on concatenated dataset of 11 nDNA regions. The same consensus trees with branch lengths untransformed is shown on the left. Maximum parsimony bootstrap values (BS_MP_), maximum likelihood bootstrap values (BS_ML_) and Bayesian posterior probabilities (PP) were denoted at internal nodes in that order; − is annotated when BS_MP_, BS_ML_ or PP_MB_ values are <50%.

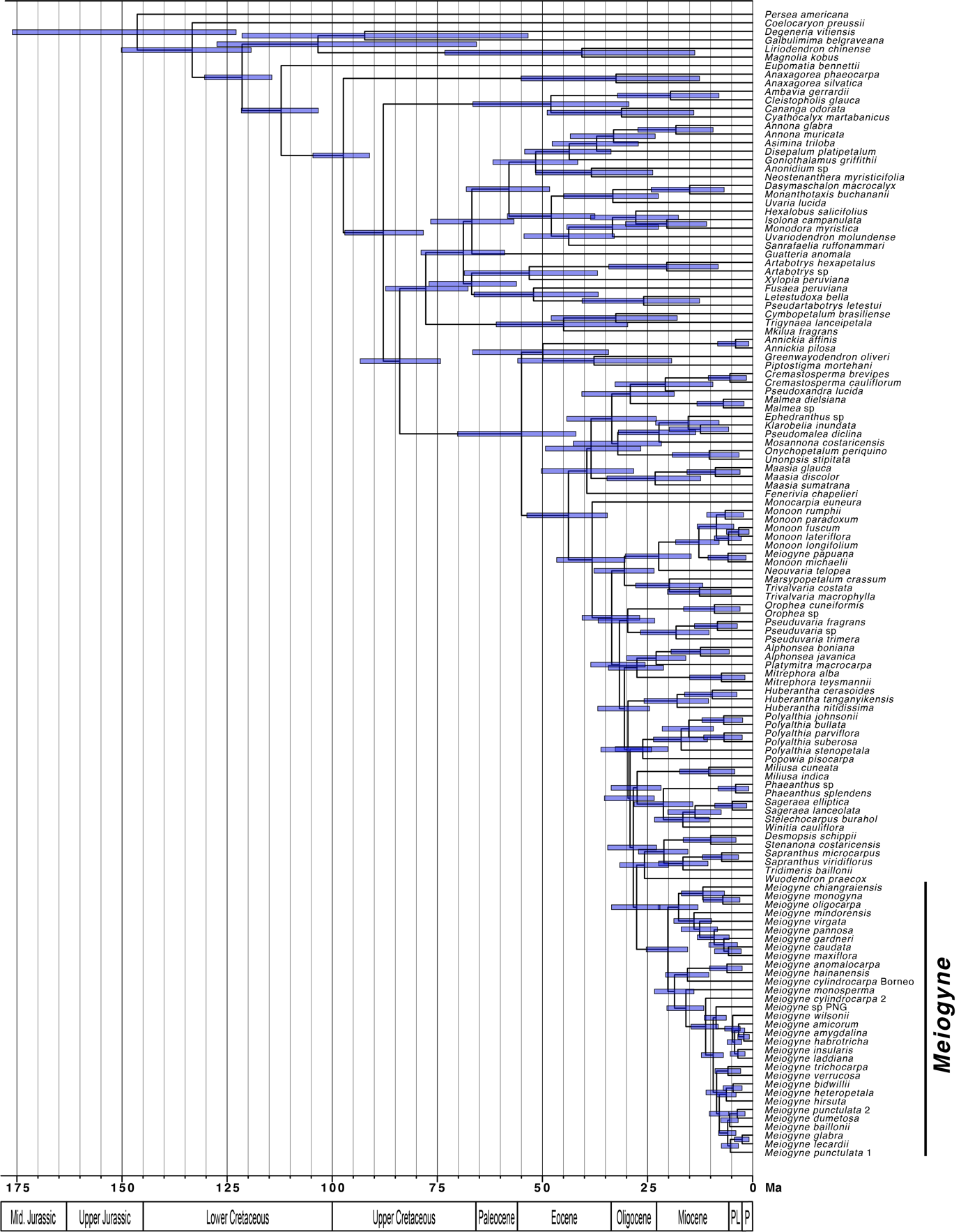

**Appendix S6** Molecular divergence time estimation. The maximum clade credibility chronogram were summarized from 10,500 post-burn-in BEAST samples. The 95% highest posterior density date ranges (95% HPDs) were indicated by blue bars.

|  | **Posterior Probability** | **Mean Divergence Time Estimate (95% HPD)** |
| --- | --- | --- |
| **Miliuseae (stem)** | 1 | 38.2 (30.4 - 46.6) |
| **Miliuseae (crown)** | 1 | 33.5 (26.9 – 40.5) |
| ***Meiogyne* (stem)** | 1 | 27.6 (22.1 – 33.6) |
| ***Meiogyne* (crown)** | 1 | 20.2 (15.5 – 25.3) |
| ***Meiogyne* Clade A (crown)** | 1 | 17.6 (13.0 – 22.5) |
| ***Meiogyne* Clade B (stem)** | 1 | 18.6 (14.0 – 23.3) |
| ***Meiogyne* Clade B (crown)** | 1 | 15.5 (10.4 – 20.7) |
| ***Meiogyne* Clade C & D (stem)** | 1 | 15.9 (11.6 – 20.4) |
| ***Meiogyne* Clade D (crown)** | 1 | 11.2 (8.2 – 14.7) |

**Appendix S7** Summary of divergence time estimates of major internal nodes from 10,500 post-burn-in BEAST samples.
